# Structural colours reflect individual quality: a meta-analysis

**DOI:** 10.1101/2020.01.01.892547

**Authors:** Thomas E. White

## Abstract

Ornamental colouration often communicates salient information to mates, and theory predicts covariance between signal expression and individual quality. This has borne out among pigment-based signals, but the potential for ‘honesty’ in structural colouration is unresolved. Here I synthesised the available evidence to test this prediction via meta-analysis and found that, overall, the expression of structurally coloured sexual signals is positively associated with individual quality. The effects varied by measure of quality, however, with body condition and immune function reliably encoded across taxa, but not age nor parasite resistance. The relationship was apparent for both the colour and brightness of signals and was slightly stronger for iridescent ornaments. These results suggest diverse pathways to the encoding and exchange of information among structural colours, while highlighting outstanding questions as to the development, visual ecology, and evolution of this striking adornment.

## Introduction

Colour is a ubiquitous channel of communication in nature and is showcased at an extreme in the service of mate choice [1,2]. A central hypothesis in evolutionary biology is that sexual selection has driven the elaboration of ornamental colouration into reliable indicators of individual quality [3], with empirical tests guided by indicator and handicap models [4,5]. These models argue that conspicuous displays are selectively favoured because their production is differentially costly (handicap) and/or constrained (index) between individuals of varying quality, and so encode honest information to potential mates. A prediction common to honesty-based models is that signals should show heightened condition-dependent expression, and the most robust support to date among ornamental colouration is found in carotenoid-based signals [6,7]. As pigments that cannot be synthesised *de novo*, all carotenoids must ultimately be acquired via diet before being incorporated into signals directly or following bioconversion. This offers ample opportunity for selection to favour mechanistic links between foraging, metabolic performance, and sexual signal expression (that is, the combined perceptual features of hue, saturation, and brightness), which is now well established, at least among birds [8,9]. Relative to our knowledge of pigment-based colouration, however, the potential for structural colours to signal individual quality remains both understudied and poorly resolved.

Unlike pigments, which are selectively absorbent, structural colours result from the selective reflectance of light by nano-structured tissues [10,11]. Accumulating evidence also suggests that the development of these structures is driven by self-assembly — such as the phase separation of keratin and cytoplasm in nascent feather barbs [12-14] — rather than the active (and ‘expensive’) cellular processes that underlie some pigmentary colour production [8]. Three general arguments have been articulated around their potential for honesty among structural colouration in sexual signalling. One is that if sufficient material is required to produce nano-architectures then it will establish a trade-off with other physiological needs that may be differentially met among individuals of varying ‘quality’, as consistent with a handicap explanation [4,6]. A non-exclusive alternative builds on the observation that features of the macro-scale expression of signals relies on the precision with which the underlying structures are organised [15,16]. If individuals heritably vary in their capacity to achieve such organisation as a result of, for example, physiological constraints on signal production, or the behavioural acquisition of stable developmental environments, then signals may serve as an index of underlying genetic quality [5]. Finally, the lack of obvious ecologically relevant material to trade-off against during signal construction, together with the self-assembly inherent in structural colours noted above, has motivated arguments against any general expectations for condition dependence *sensu lato* [12]. Though experimental work is able to partition these hypotheses in some contexts [17], most empirical studies to date have focused on the overarching question of honesty by examining the predicted covariance between fitness-related traits and signal expression. This has provided valuable insight into the central question, but diversity in signal designs, measures of ‘quality’, and taxonomy have presented a challenge for qualitative synthesis. Modern quantitative methods, however, provide robust tools for controlling for and capitalising on such variation (e.g via mixed-effects models and meta-regression), and so can offer substantive answers to longstanding questions [18].

Here I used phylogenetically controlled meta-analysis and meta-regression to examine whether structural colour signals encode salient information on individual quality. Specifically, I synthesised estimates of correlations between measures of individual quality and signal expression to test the prediction of condition dependence, before examining methodological and theoretically derived mediators of effect-size variation among studies.

## Methods

### Literature search and study selection

I conducted a systematic literature search using *Web of Knowledge* and *Scopus* databases for publications up to September 2019, using the query ((colour OR color OR pigment) AND signal AND (quality OR condition OR condition dependent OR condition dependence OR ornament) OR honest*), as well as searching the references of included texts. This produced 3482 unique studies, from which 41 were ultimately suitable for quantitative synthesis following the screening of titles and abstracts (where n = 3430 were excluded for clear irrelevance), and full texts (see Fig. S1 for PRISMA statement). I used the R package ‘revtools’ v0.4.1 for title and abstract screening [19]. I included all experimental and observational studies that quantified the relationship between intersexual structural colour signal expression (via the measurement of hue, saturation, or brightness, or a composite thereof) and any one of age, body condition (size, size-corrected mass, or growth rate), immune function (oxidative damage, PHA response, circulating CORT or testosterone) or parasite resistance as a measure of individual quality. I excluded studies that conflated the structural and pigmentary contributions to signal expression during measurement or manipulation, only studied sexually immature juveniles, focused exclusively on intrasexual signalling, used human-subjective assessments of colouration (such as colour swatches or viewer rankings), or which did not provide adequate data in the form of raw effect sizes, or summary statistics or figures from which effect sizes might be estimated.

### Effect size calculation

I used the correlation coefficient, Pearson’s r, transformed to Fisher’s z (given its preferable normalizing and variance-stabilizing qualities) as the effect size describing the relationship between colouration and individual quality for meta-analysis. These effects were extracted directly from text or figures, using the R package ‘metadigitise’ v1.0 [20], where possible (n = 102), or was otherwise converted from available test statistics or summary data (n = 84).

### Meta-analyses

I ran both phylogenetic multi-level meta-analytic (intercept-only, MLM) and multi-level meta-regression (MLMR) models, using the package ‘metafor’ v2.1-0 [21] in R v3.5.2 [22]. Almost all studies reported multiple effects through the estimation of several colour metrics or multiple measures of individual quality, so I included both a study- and observation-level random effect in all models. From my MLM model I estimated a meta-analytic mean (i.e., intercept) effect size, which describes the overall support for the honesty of structural colour signals. I accounted for phylogenetic non-independence between effect sizes in all models by estimating relationships among species using the Open Tree of Life database [23], accessed via the R package ‘rotl’ v3.0.10 [24]. Given the resulting tree topology, I estimated a correlation matrix from branch lengths derived using Grafen’s method [25] assuming node heights raised to the power of 0.5. Though this does not account for evolutionary divergence, it grants an approximate estimate of relatedness by accounting for phylogenetic topology (Fig. S2).

I then used separate MLMR models to examine the effects of moderators, both theoretical and methodological, which may be expected to alter the strength of the signal/quality relationship. These included the measure of individual quality used—body condition, age, immune function, or parasite resistance (as defined above)—since ‘quality’ is multivariate (discussed below). There is a suite of metrics available for measuring colour, though they typically center on quantifying the perceptually relevant features of hue (the unique colour), saturation (spectral purity), and brightness, or a composite thereof [26]. I therefore classified every measure as such in order to test which, if any, signal features contain salient information on mate quality. In broad terms, the greater nano-structural organisation and/or material required to generate more saturated and (to a lesser extent) brighter signals predicts a positive correlation between these features and individual quality. Signal hue, by contrast, is a directionless measure in the sense that there is no clear biophysical reason for predicting consistent among-individual shifts toward longer or shorter wavelengths as a function of individual quality across taxa, and so I ignored the sign of correlations for estimates of hue alone. I also tested the effect of signal iridescence (i.e. the presence/absence of iridescent colouration), which I coded according to information presented in-text or via secondary sources (including personal observation). The rationale was twofold. For one, all iridescence arises from coherent light-scattering [27]. All things being equal, coherent light-scatterers demand a level of architectural organisation beyond that of incoherent scatterers (i.e. white colours) and possibly non-iridescent colours too, and so offers an indirect test of the hypothesised link between the demands of nano-scale organisation and signal honesty [14,16]. Second, iridescence is an inherently temporal feature of visual communication which may provide an additional or alternate conduit of information to potential mates, above and beyond that which is possible using non-iridescent signals (17,28,29; though this possibility remains unexplored directly). In both cases the prediction is a stronger correlation between colouration and quality among iridescent, as opposed to non-iridescent, ornaments. Finally, and following the above, I secondarily examined the effects of both quality measures and colour metrics within each of the four taxonomic classes represented across the literature. I focused on these two moderators alone because potential taxonomic variation in the mechanistic links between colouration and individual quality are most likely to manifest via these moderators, and because the limited available data precludes the testing of all moderators on a per-class basis (note that even within these moderators, estimates were not possible across all taxonomic groups).

With respect to methodology I considered study type, given my inclusion of both experimental and observational studies, as well as the sex of focal animals. I also coded whether studies included measurements of non-sexual traits as controls in tests of *heightened* condition-dependence (see discussion). The prediction being that that studies including non-sexual controls may report larger effects than those without, given that many traits will scale with condition to some extent [30]. Note that both signal iridescence and the presence of controls were coded as binary (0/1) for simplicity in testing their respective predictions.

### Publication bias

I explored evidence for publication bias by visually inspecting funnel plots of effect sizes versus standard errors (Fig. S3) and using an Egger’s test on an intercept-only MLMA that included the random effects described above [31].

### Data availability

All data and code are available via GitHub (https://github.com/EaSElab-18/ms_metacol) and are persistently archived through Zenodo (https://dx.doi.org/10.5281/zenodo.3718617).

## Results

The final dataset comprised 186 effect sizes, across 28 species, from 41 studies [6, 17, 32-71]. Of those 186 effects, 117 were drawn from birds, 22 from insects, 28 from reptiles, and 11 from arachnids (Table S1; Fig. S2). As predicted, I found a positive overall correlation (i.e. meta-analytic mean effect) between individual quality and structural colour signal expression (*Z* = 0.1573, 95% CI = 0.084 to 0.231; Fig. 1; Table 1). The heterogeneity of effect sizes — that is, the variation in effect size estimates after accounting for sampling error — was high (I^2^ = 80.42%, 95% CI = 77.26 to 83.01), as is typical of meta-analytic data in ecology and evolutionary biology [72]. A small amount of heterogeneity was explained by among-study effects (I^2^ = 14.21%, 95% CI = 8.97 to 20.20), and only a very weak phylogenetic signal was evident (I^2^ = 2.17%, 95% CI = 1.18 to 3.47).

**Table 1:** Full parameter estimates from MLM and MLMR models of the mediators of the correlation between structural colour signal expression and individual quality. Shown are sample sizes, mean Fisher’s z values and lower and upper 95% confidence intervals, and heterogeneity. Estimates whose 95% confidence intervals do not overlap zero are indicated in bold. Note that iridescence and the inclusion of controls are coded as binary (0/1), and so represent a test of difference in effect-sizes between their counterpart categories (see main text).

| Model | n | Mean (Zr) | Lower CI | Upper CI | I<sup>2</sup> (%) |
| --- | --- | --- | --- | --- | --- |
| Overall (intercept-only) | 186 | **0.157** | **0.084** | **0.231** | 80.42 |
| Quality measure |  |  |  |  | 79.96 |
| age | 37 | 0.015 | -0.119 | 0.148 |  |
| body condition | 102 | **0.190** | **0.099** | **0.282** |  |
| immune function | 11 | **0.353** | **0.126** | **0.580** |  |
| parasite resistance | 36 | 0.114 | -0.023 | 0.252 |  |
| Component |  |  |  |  | 80.32 |
| hue | 50 | **0.224** | **0.123** | **0.345** |  |
| saturation | 57 | 0.076 | -0.029 | 0.181 |  |
| brightness | 60 | **0.171** | **0.079** | **0.264** |  |
| composite | 19 | 0.120 | -0.056 | 0.296 |  |
| Sex |  |  |  |  | 80.78 |
| female | 29 | 0.131 | -0.004 | 0.267 |  |
| male | 146 | **0.161** | **0.080** | **0.241** |  |
| not distinguished | 11 | 0.183 | -0.046 | 0.413 |  |
| Study type |  |  |  |  | 80.39 |
| experimental | 52 | **0.216** | **0.104** | **0.329** |  |
| observational | 134 | **0.128** | **0.058** | **0.199** |  |
| Optics |  |  |  |  | 82.64 |
| iridescent (vs not) | 67 | **0.150** | **0.009** | **0.291** |  |
| Control |  |  |  |  |  |
| included (vs not) | 28 | 0.071 | -0.117 | 0.260 | 83.28 |

**Figure 1:**
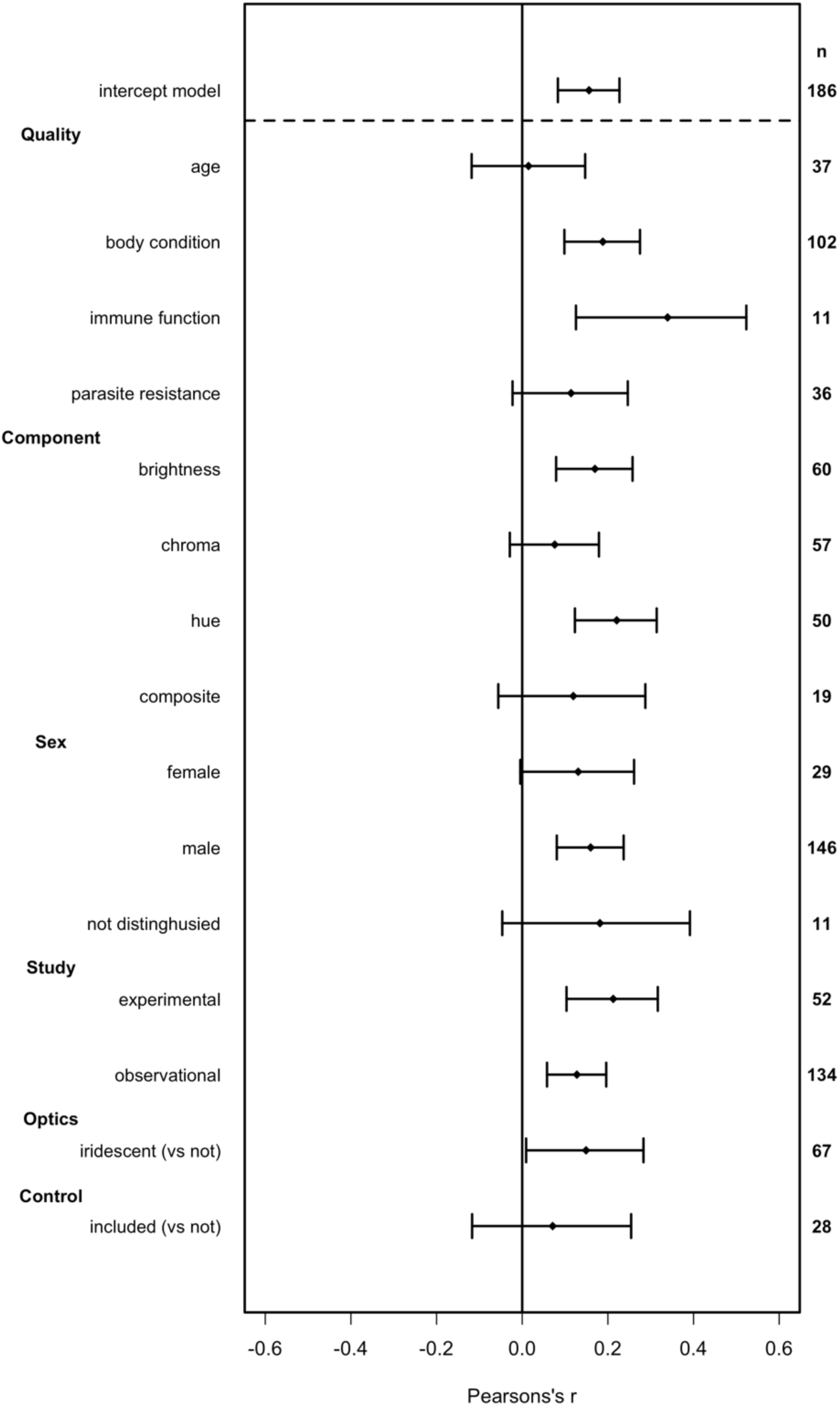
Forest plot of the mediators of the correlation between structural colour signal expression and individual quality. Shown are Pearson’s correlations back transformed from Fisher’s z, with 95% confidence intervals about the mean. Sample sizes are displayed on the right. ‘Composite’ refers to measures that conflate hue, saturation, and brightness (such as PCA), while ‘not distinguished’ refers to studies in which the sex of focal animals was either not specified, or males and females were pooled. Note that iridescence and the inclusion of controls are coded as binary (0/1), and so represent a test of difference in effect-sizes between their counterpart categories (see main text).

Of the measures of quality considered, body condition and immune function were reliably positively correlated with structural colour expression, while age and parasite resistance were not (see Table 1 for all corresponding numerical results henceforth). This varied slightly across taxa, however, with a robust effect of condition on colouration apparent across all groups, while effects of age and parasite resistance we apparent among insects and bird, respectively (table S2; though these estimates are based on limited within- and between-taxa samples). Both the hue and brightness of signals were similarly informative channels on-whole, while chromaticity was not consistently associated with individual quality across taxa (though this varied by taxa; table S2), nor was any correlation apparent when composite measures of colouration were used. Iridescent signals were subject to slightly stronger positive correlations than non-iridescent signals across all measures of condition. Signal honesty was apparent among males only though the weak, borderline effect and much smaller sample among females (n = 29/186 versus 146/186 for males) suggests a male bias in the literature similar to that in related fields [73], which may have partly driven this outcome. Experimental studies tended to report marginally stronger correlations than observational assays, which most likely reflects slightly exaggerated experimental manipulations of condition relative to natural variation [30]. Finally, the majority of studies (n = 36) did not include measurements of non-sexual control traits in tests of *heightened* condition dependence, though I found no clear difference in effect-size estimates between those that did and did not.

### Publication bias

Visual inspection of the funnel plot showed little asymmetry (Fig. S3), as supported by non-significant Egger’s tests (t_184_ = −0.5535, p = 0.5806), which suggests a minimal influence of missing data on effect size estimates.

## Discussion

Ornamental colouration may be a reliable conduits of information on mate quality, though evidence for the predicted covariance between signal expression and mate quality among structural, as opposed to pigmentary, signals is equivocal. Here I found meta-analytic support for this link in the form of a positive correlation between structural colour expression and individual quality (Fig. 1), consistent with honesty-based models of sexual signal evolution [4,5]. The strength of the overall correlation, though moderate [74], was commensurate with meta-analytic estimates from pigment-based sexual signals [8,9,75], and suggests that structural colouration may similarly serve a reliable indicator of individual quality.

Quality is a multivariate feature of individuals, and this is reflected in the effect-size variation between measures. Both condition (as narrowly defined above), and proxy measures of immune system integrity were on-average positively correlated with signal expression across all taxa in which those relationships have been examined. This is consistent with experimental work showing that body mass and immune function are responsive to ecologically salient stressors, with consequences for colour production. Among birds, for example, disease and dietary stress produce abnormalities in the keratin barbules that contribute to colouration [16,76,77], while in butterflies the organisation of wing-scale architectures is disrupted by nutritional and environmental stress during pupal (hence, wing-structure) development [36,78]. In contrast, neither age nor parasite resistance were consistently informative of mate quality, though this varied slightly across taxa. These latter measures are often predicated on, or susceptible to, the mechanical degradation of structures post-development. Thus, the inherently heightened variability of sexual signals combined with parasite-induced damage (ectoparasite, in particular) and/or accumulated wear with age, combined with varied mechanisms of colour production of across animal classes, may compound to render the signals less accurate predictors on balance [59,79,80]. Curiously, the near inverse relationship was recently identified in a meta-analysis of carotenoid-based signalling. Weaver et al. [8] examined correlations across similar categories of quality as those used here but found no consistent relationship between signals and either of body condition or immune function. Given the fundamental optical and developmental differences between structural and pigmentary colour production (described above) the potential exists for each to signal unique aspects of individual mate quality, as is suggested by the totality of this work. This has also been directly supported by limited empirical work [65] and may hold more broadly as an explanation for the often-integrated use of structural and pigmentary mechanisms in sexual colouration.

Colour is often assumed to be the central conduit of information exchange given its relative stability under variable natural illumination [81,82], though my results suggest both the colour and brightness of signals are similarly informative, considering the evidence to date (Fig. 1; Table S2). Furthermore, I identified slightly stronger condition dependence among iridescent, as opposed to non-iridescent, patches. While the underlying architecture varies across taxa, all iridescent colouration arises from coherent light interference and so may demand a level of architectural organisation beyond that of incoherent scattering [11,27], as well as non-iridescent coherent scattering (though evidence for the latter possibility is limited; [14]). Iridescence also introduces temporal structure to signals since the colour appearance depends on the precise arrangement of signals, viewers, and light sources. These combined features may render iridescent colouration particularly suitable as bearers of information [29] and so contribute to the ubiquity of the phenomenon [83,84]. Though only indirectly considered here, as few studies quantify between-individual variation in iridescence itself, this idea has found more immediate support via condition-dependent variation in signal angularity [17], and a predictive relationship between iridescence and mating success [85]. Empirically unravelling the function and perceptual significance of iridescence in the context of sexual signalling—where the effect is seen at its most extreme—remains an active challenge [28]. More generally, these results affirm the view that the extended spectral and temporal repertoire available to structural colours may facilitate the exploration of distinct ‘signalling niches’, with tangible evolutionary consequences [1,54].

By integrating the development of signal structure and fitness-related traits, structural colours may serve as informative signals during mate choice. A holistic understanding, however, awaits progress on several fronts. Most significant is the inclusion of appropriate non-sexual controls. Given that many traits will scale with overall condition, the ultimate evidence for handicap models lies in the demonstration of *heightened* condition-dependence among sexual traits. Though I found no clear difference in effect size estimates between studies with and without such controls the small sample size was limiting, and moreover represents a conceptual limitation that remains pervasive [30]. Partitioning indicator and handicap models of signal evolution and understanding the nature of direct and/or indirect benefits being signalled, are key challenges which requires both experimental and quantitative-genetic study [17]. The development of structural colours during ontogeny is also a central front for progress, and studies among invertebrates (which offer benefits in terms of tractability) would be invaluable in complementing the excellent work accumulating on birds [12-14]. Finally, signalling ecology should remain front-of-mind as accumulating evidence, consistent with that presented here, continues to highlight the inherent spatio-temporal complexity of signals and visual systems [86-88]. This offers exciting opportunities for integrative studies of signal development, production, and perception, which will fuel a richer view of this pervasive adornment of the natural world.

## Acknowledgments

I’m grateful to Daniel Noble and Alistair Senior for advice on meta-analysis, and Elizabeth Mulvenna and Cormac White for their endless support. I appreciate the efforts of three anonymous reviewers whose thoughtful suggestions considerably improved this study.

