## Supplementary material for "Structural colours reflect individual quality: a meta-analysis"

### Supplementary material for White TE (2020) Structural colours reflect individual quality: a meta-analysis.

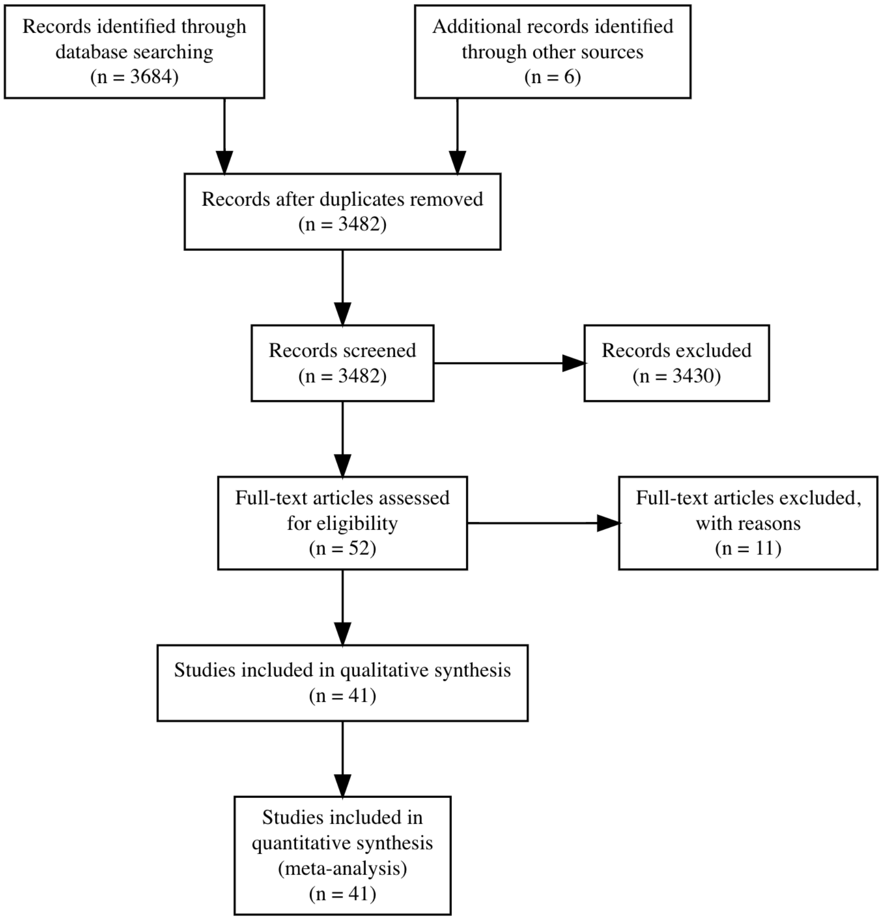


Figure 1: PRISMA diagram depicting the systematic search strategy for literature testing the relationship between the expression of structural colour signals and individual quality.

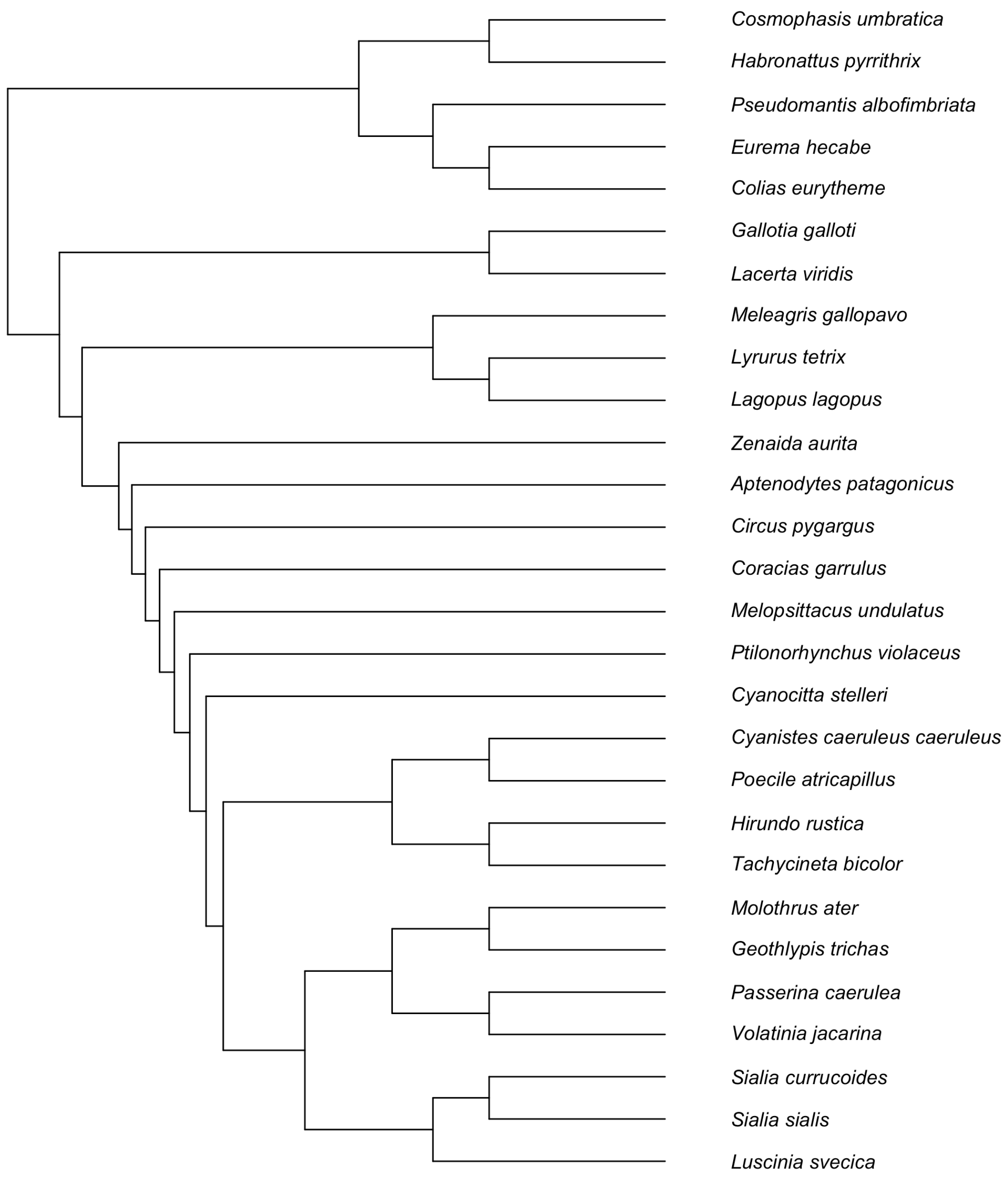


Figure 2: Phylogenetic tree of the taxa included in this study.

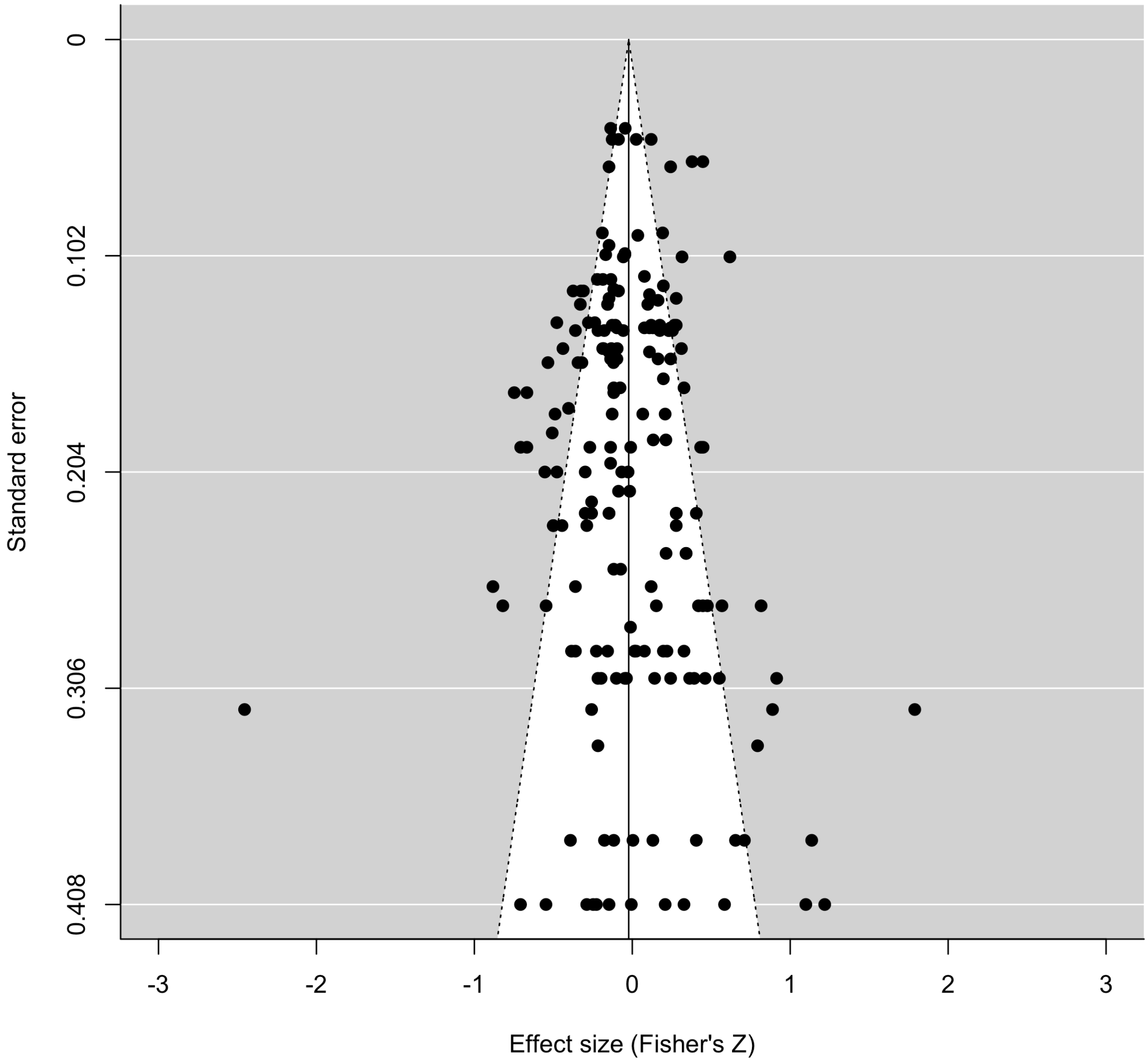


Figure 3: Funnel plots of effect sizes against their standard error, with 95% pseudo confidence interval denoted by dashed lines.

**Table S1:** Summary of the included studies and key data therein, including the class and species of the study organism, the sex of individuals examined, the colour metrics used, as well as measure of quality considered (see main text for category and colour definitions).

| Study | Year | Class | Species | Sex | Colour | Quality |
|---|---|---|---|---|---|---|
| Bajer et al. | 2012 | Reptilia | *Lacerta viridis* | m | chroma, brightness | condition |
| Balenger et al. | 2007 | Aves | *Sialia currucoides* | m, f | composite | condition |
| Barry et al. | 2015 | Insecta | *Pseudomantis albofimbriata* | f | brightness | condition |
| Beck et al. | 2015 | Aves | *Tachycineta bicolor* | m, f | composite | parasite |
| Bitton et al. | 2008 | Aves | *Tachycineta bicolor* | m | hue, chroma, brightness | age |
| Bitton et al. | 2008 | Aves | *Tachycineta bicolor* | m, f | brightness | age |
| D'Alba et al. | 2011 | Aves | *Poecile atricapillus* | m, f | brightness, chroma | parasite |
| Doucet | 2002 | Aves | *Volatinia jacarina* | m | brightness | condition |
| Doucet & Montgomerie | 2003 | Aves | *Ptilonorhynchus violaceus* | m | brightness | condition, parasite |
| Freeman-Gallant et al. | 2010 | Aves | *Geothlypis trichas* | m | brightness | condition |
| Griggio et al. | 2010 | Aves | *Melopsittacus undulatus* | m | hue, chroma, brightness | immune |
| Grindstaff et al. | 2012 | Aves | *Sialia sialis* | m, f | composite | condition, immune |
| Henderson et al. | 2013 | Aves | *Cyanistes caeruleus* | f | chroma | condition |
| Hill et al. | 2005 | Aves | *Meleagris gallopavo* | m | composite | parasite |
| Kemp et al. | 2006 | Insecta | *Colias eurytheme* | m | hue, brightness | condition |
| Kemp & Rutowski | 2007 | Insecta | *Eurema hecabe* | m | hue, brightness | condition |
| Kemp | 2006 | Insecta | *Colias eurytheme* | m | hue, chroma, brightness | age |
| Kemp | 2008 | Insecta | *Eurema hecabe* | m | hue, brightness | condition |
| Keyser & Hill | 2000 | Aves | *Passerina caerulea* | m | hue, chroma, brightness | |
| Keyser & Hill | 1999 | Aves | *Passerina caerulea* | m | hue, chroma, brightness | condition |
| Lim & Li | 2007 | Arachnida | *Cosmophasis umbratica* | m | hue, brightness | age, condition |
| McGraw et al. | 2002 | Aves | *Molothrus ater* | m | hue, chroma, brightness | condition |
| Megia-Palma et al. | 2016 | Reptilia | *Gallotia galloti* | m, f | hue, chroma, brightness | parasite |
| Molnar et al. | 2012 | Reptilia | *Lacerta viridis* | m, f | brightness, chroma | condition, parasite |
| Molnar et al. | 2013 | Reptilia | *Lacerta viridis* | m | chroma, brightness | parasite |
| Mougeot et al. | 2006 | Aves | *Circus pygargus* | m | hue, chroma | condition |
| Mougeot et al. | 2005 | Aves | *Lagopus lagopus* | m, f | hue, brightness | age, parasite |

| | | | | | | |
|---|---|---|---|---|---|---|
| Nicolaus et al. | 2007 | Aves | *Aptenodytes patagonicus* | m | hue, chroma, brightness | age |
| Perrier et al. | 2002 | Aves | *Hirundo rustica* | m, f | chroma | age |
| Peters et al. | 2006 | Aves | *Cyanistes caeruleus* | m | chroma | immune |
| Peters et al. | 2011 | Aves | *Cyanistes caeruleus* | m, f | hue, chroma, brightness | condition |
| Quinard et al. | 2017 | Aves | *Zenaida aurita* | m | brightness | condition |
| Roberts et al. | 2009 | Aves | *Cyanistes caeruleus* | m, f | chroma | immune |
| Schull et al. | 2016 | Aves | *Aptenodytes patagonicus* | m, f | hue, chroma, brightness | age, parasite |
| Siefferman et al. | 2005 | Aves | *Sialia sialis* | m | composite | age |
| Siitari et al. | 2007 | Aves | *Tetrao tetrix* | m | chroma, brightness | immune |
| Silva et al. | 2008 | Aves | *Coracias garrulus* | m, f | composite | condition |
| Smiseth et al. | 2001 | Aves | *Luscinia svecica* | m | hue, chroma, brightness | age, condition |
| Taylor et al. | 2011 | Arachnida | *Habronattus pyrrithrix* | m | hue, chroma, brightness | condition |
| Viblanc et al. | 2016 | Aves | *Aptenodytes patagonicus* | m, f | hue, brightness | condition, immune |
| Zirpoli et al. | 2013 | Aves | *Cyanocitta stelleri* | m | hue, chroma, brightness | condition, parasite |

**Table S2:** Parameter estimates from MLMR models of two mediators of the correlation between structural colour signal expression and individual quality within each class of organism studied. Shown are sample sizes and 95% confidence intervals (lower bound, upper bound) for Fisher's z values. Estimates whose 95% confidence intervals do not overlap zero are indicated in bold. NA's indicate contrasts for which no data are available.

| | *Aves* | | *Insecta* | | *Arachnida* | | *Reptilia* | |
|---|---|---|---|---|---|---|---|---|
| **Model** | **n** | **Zr (CI)** | **n** | **Zr (CI)** | **n** | **Zr (CI)** | **n** | **Zr (CI)** |
| Overall | 117 | **0.034, 0.292** | 22 | **0.026, 0.495** | 19 | -0.128, 0.439 | 28 | -0.163, 0.341 |
| Quality | | | | | | | | |
| age | 30 | -0.183, 0.126 | 3 | **0.191, 0.708** | 4 | -0.264, 0.233 | 0 | NA |
| body condition | 54 | **0.014, 0.239** | 19 | **0.105, 0.305** | 15 | **0.103, 0.419** | 14 | **0.092, 0.477** |
| immune function | 11 | **0.102, 0.582** | 0 | NA | 0 | NA | 0 | NA |
| parasite resistance | 22 | **0.117, 0.477** | 0 | NA | 0 | NA | 14 | -0.277, 0.109 |
| Component | | | | | | | | |
| hue | 37 | **0.166, 0.503** | 10 | -0.034, 0.239 | 8 | **0.116, 0.402** | 8 | -0.032, 0.521 |
| saturation | 37 | -0.038, 0.273 | 1 | **0.032, 0.851** | 4 | **0.065, 0.309** | 15 | -0.206, 0.199 |
| brightness | 24 | **0.001, 0.274** | 11 | **0.225, 0.486** | 7 | -0.099, 0.244 | 5 | -0.153, 0.517 |
| composite | 19 | -0.104, 0.338 | 0 | NA | 0 | NA | 0 | NA |
